# Common recruitment of angular gyrus in episodic autobiographical memory and bodily self-consciousness

**DOI:** 10.1101/345991

**Authors:** Lucie Bréchet, Petr Grivaz, Baptiste Gauthier, Olaf Blanke

## Abstract

Parietal cortex and adjacent parts of the temporal cortex have recently been involved in bodily self-consciousness (BSC) and in episodic autobiographical memory (EAM). However, the neuroanatomical relationship between both fundamental aspects of self-related processing remains currently unexplored. Here we investigated whether regions in the inferior parietal lobule (IPL) that have been involved in BSC (self-location and first-person perspective) are also activate in studies investigating autobiographical memory. To examine this relation, we performed a meta-analytical study based on functional neuroimaging studies on EAM and SAM and compared them with BSC activations. We report an anatomical overlap bilaterally in the angular gyrus (AG), but not in other parietal or temporal lobe structures between BSC and EAM. Moreover, there was no overlap between BSC and SAM, suggesting that the bilateral AG is a key structure for the conscious re-experiencing of past life episodes (EAM) and the conscious on-line experience of being located and experiencing the world in first-person (BSC).

## 1. Introduction

The subjective feeling of a self that is experienced as residing in one’s body, which is localized at a specific position and from where I perceive the world has been defined as bodily self-consciousness (BSC) (Blanke and Metzinger, 2009; Blanke et al., 2015). Although the experimental investigation of BSC and the underlying brain networks remains a challenge, recent advances in digital technologies have provided compelling ways of successfully inducing illusory states of BSC in healthy individuals by manipulating multisensory bodily cues (Ehrsson, 2007; Lenggenhager et al., 2007; Petkova and Ehrsson, 2008). For example, illusory self-location and first-person perspective were manipulated in a fMRI study (Ionta et al., 2011) by applying tactile stroking to the back of a participant, while simultaneously displaying the stroking on the back of a virtual body via a head mounted display. These experimentally-induced changes in BSC have been linked to activity at the temporo-parietal junction (TPJ; Brodmann area 39/40). Likewise, lesions, seizures or electrical brain stimulation at the TPJ also result in changes in self-location and first-person perspective (Blanke et al., 2002, 2004; Ionta et al., 2011; De Ridder et al., 2007). Thus, patients suffering from so-called out of body experiences (OBE) caused by brain damage at the TPJ, subjectively experience the world from an illusory disembodied self-location with an inverted direction of first-person perspective and they self-identify with this elevated position (Blanke and Arzy, 2005; Heydrich and Blanke, 2013; Ionta et al., 2011).

The subjective sense of self in time that enables us to re-experience ourselves in the past and mentally project ourselves into the future, i.e. autonoetic consciousness, has been considered the defining aspect of episodic autobiographical memory (EAM) recollection (Tulving, 1985a). In addition, Tulving proposed that, contrary to EAM, semantic autobiographical memory (SAM) is characterized by noetic consciousness, i.e. the general self-awareness of personal facts, independent of re-experiencing particular life episodes. Traditionally, since the description of severely amnesic patients with damage to the medial temporal lobe (MTL) (Scoville, 1957; Steinvorth et al., 2005), many studies, including neuroimaging studies, confirmed the essential role of MTL structures in memory (Cabeza et al., 2007; Clark and Maguire, 2016; Daselaar et al., 2008; St. Jacques et al., 2015; Svoboda et al., 2006). More recently, neuroimaging studies of EAM retrieval (Bellana et al., 2017; Cabeza and St Jacques, 2007; Gilmore et al., 2015; Rugg and Vilberg, 2013; Rutishauser et al., 2017; Sestieri et al., 2011, 2017; St Jacques and Schacter, 2013; Uncapher and Wagner, 2009; Wagner et al., 2005) confirmed the role of MTL structures, but also pointed out persistent and robust activations in the inferior parietal lobule (IPL), particularly the angular gyrus (AG; Brodmann area 39). Interestingly, patients with lateral parietal lesions oerform normally in objective EAM tasks, however, it has been found that the vividness, richness and subjective confidence in experiencing their personal memories is diminished (Ben-Zvi et al., 2015; Berryhill et al., 2007; Hower et al., 2014; Rugg and King, 2017; Simons et al., 2010). These studies suggest that the sole engagement of MTL structures may not be sufficient for the full multimodal and conscious experiences, which accompany subjective, detail-rich and self-related EAM recollection, and that additional regions, such as those in the IPL, are involved as well.

Despite several reviews discussing the role of parietal lobe in BSC (Blanke, 2012; Blanke et al., 2015; Serino et al., 2013) as well as the recent interest in the contribution of parietal lobe to EAM (Igelström and Graziano, 2017; Moscovitch et al., 2016), it is currently unknown whether and to what extent BSC and autobiographical memories (whether episodic or semantic) engage the same or distinct brain regions, especially in the IPL. In the current study, we performed a systematic quantitative coordinate based meta-analysis (Eickhoff et al., 2009, 2012) on human functional neuroimaging studies in order to examine whether and where brain regions associated with autobiographical memories share common or distinct neural substrates. This was done separately for EAM and SAM and compared with BSC-related activations (related to changes in self-location and first-person perspective) as reported by Ionta et al. (2011). Based on the evidence that both EAM and BSC recruit IPL, we hypothesized that there would be an anatomical overlap in this structure between BSC and autobiographical memory, specifically for the episodic (EAM), but not the semantic aspects (SAM).

## 2. Methods

### 2.1. Selection of studies and inclusion criteria for EAM and SAM meta-analyses

For both the episodic as well as the semantic autobiographical memory studies, we conducted a comprehensive and systematic search of the literature using PubMed <u>(www.pubmed.org)</u> and Web of Knowledge <u>(www.webofknoledge.org).</u> The following combination of keywords was used for episodic autobiographical memory: “autobiographical memory”, “episodic”. For the semantic autobiographical memory, we selected: “autobiographical memory”, “semantic”. The reference lists of the included studies and several previous meta-analyses were used to find studies (Kim, 2016; Martinelli et al., 2013; Svoboda et al., 2006). Studies using positron emission tomography (PET) or functional magnetic resonance imaging (fMRI) were included in the analyses. Studies were considered only if activation coordinates were reported in standardized Montreal Neurological Institute (MNI) or Talairach (TAL) coordinates and the analysis reported on the whole brain. All Talairach coordinates were transformed into the MNI coordinates using a linear transformation (Lancaster et al., 2007).

Only study results in healthy subjects with no neurological or psychiatric disorders, brain lesions or pharmacological manipulations were considered. Studies including both younger and older healthy participants were included in the analysis. No single subject studies were considered for the analyses. If articles reported several experiments with independent samples, then these experiments were considered individually in the analysis. Visual as well as auditory cues were included irrespectively of their emotional valence. Both recent and remote memories for the EAM analysis were included. Personal events and judgments of the self-versus others were included in the SAM analysis.

#### 2.1.1. Episodic autobiographical memory (EAM) studies

Forty-one experiments investigated EAM **(Supplementary Table 1)** and this meta-analysis included 588 foci and 813 subjects. We focused primarily on studies investigating self-related, personally-relevant EAM irrespective of their specificity, age or control task. Thus, we included both recent and remote EAM (e.g., Addis et al., 2012; Oddo et al., 2010), using semantic memory (Donix et al., 2010; Holland et al., 2011) as well as low level baseline condition (e.g., rest or pseudo words) (Nadel and Moscovitch, 1997; Piolino et al., 2008) as control tasks.

**Table 1.**
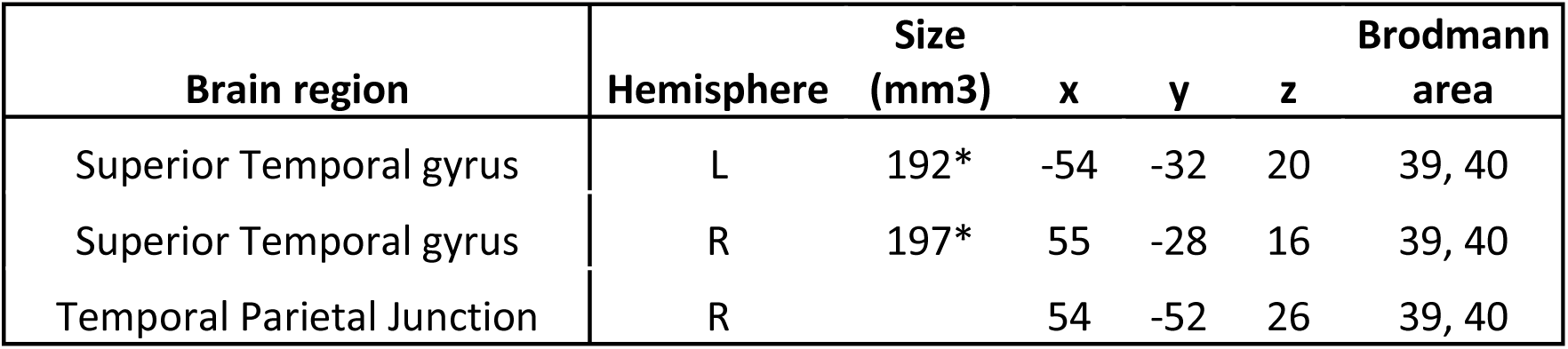
Results from BSC experimental study (N=22) FDR-corrected, p < 0.05 and neurological patients study (N=9) based on voxel-based lesion symptom mapping (VLSM) FDR-corrected, p < 0.01. R= right, L= left, B= bilateral; x, y, z are the coordinates in MNI space. Label provided using the Brodmann atlas *active cluster size in voxels

#### 2.1.2. Semantic autobiographical memory (SAM) studies

We selected twenty-five experiments examining SAM **(Supplementary Table 2)** and included 314 foci and 396 subjects. Particularly, we included studies investigating familiar, self-relevant semantic information (e.g., faces, places, objects), but also self-trait judgements (Sugiura et al., 2009, 2011). Control task in the SAM category included unfamiliar information or other-trait judgements (e.g., Gutchess et al., 2007; Jenkins et al., 2008).

**Table 2.**
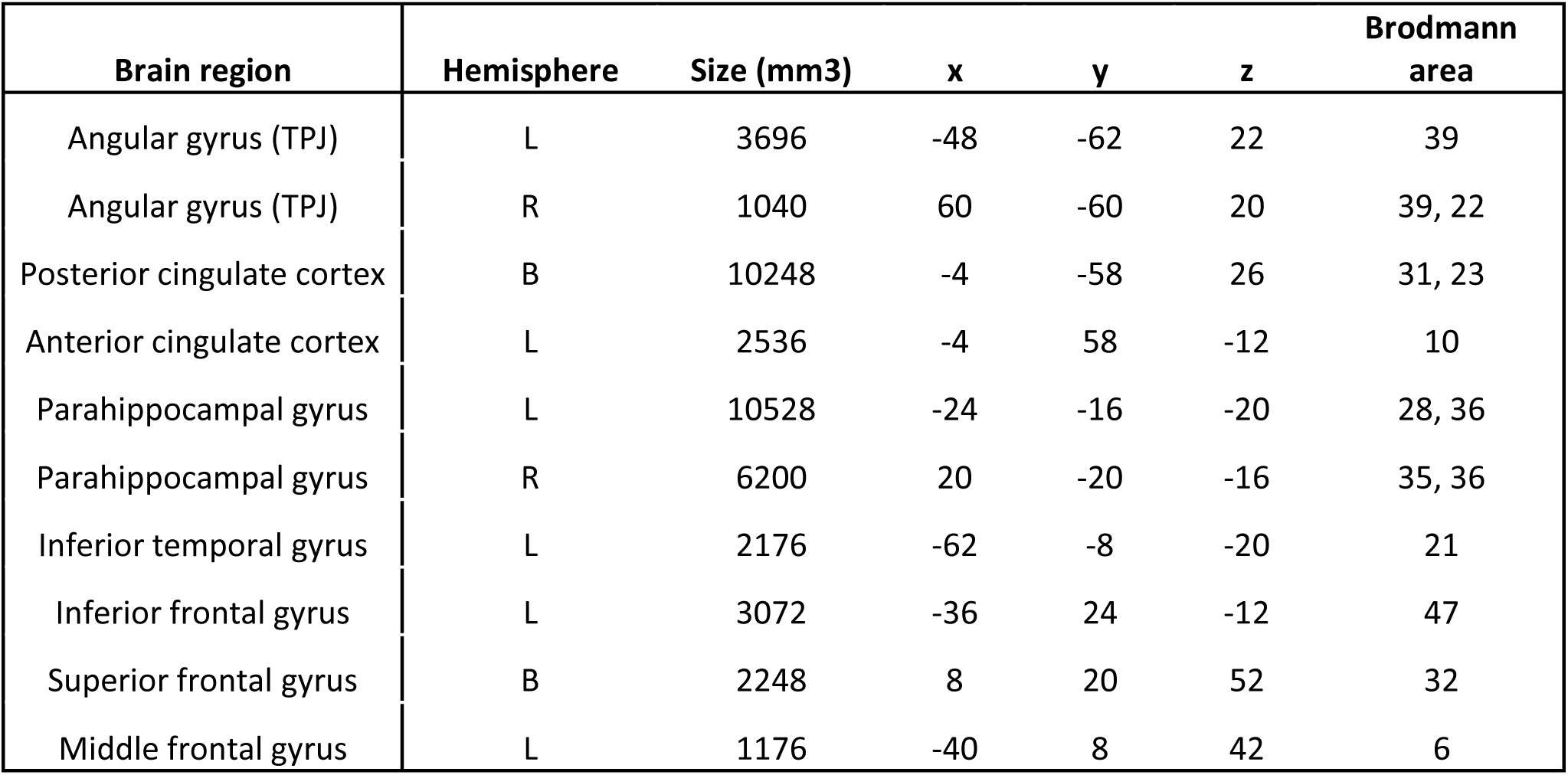
Results from EAM meta-analysis. Within-cluster FEW-corrected p < 0.05 with p < 0.001 (uncorrected) as the cluster forming threshold. R= right, L= left, B= bilateral; x, y, z are the coordinates in MNI space. Label provided using the Brodmann atlas *active cluster size in voxels

| Brain region | Hemisphere | Size (mm3) | x | y | z | Brodmann area |
| --- | --- | --- | --- | --- | --- | --- |
| Angular gyrus (TPJ) | L | 3696 | -48 | -62 | 22 | 39 |
| Angular gyrus (TPJ) | R | 1040 | 60 | -60 | 20 | 39, 22 |
| Posterior cingulate cortex | B | 10248 | -4 | -58 | 26 | 31, 23 |
| Anterior cingulate cortex | L | 2536 | -4 | 58 | -12 | 10 |
| Parahippocampal gyrus | L | 10528 | -24 | -16 | -20 | 28, 36 |
| Parahippocampal gyrus | R | 6200 | 20 | -20 | -16 | 35, 36 |
| Inferior temporal gyrus | L | 2176 | -62 | -8 | -20 | 21 |
| Inferior frontal gyrus | L | 3072 | -36 | 24 | -12 | 47 |
| Superior frontal gyrus | B | 2248 | 8 | 20 | 52 | 32 |
| Middle frontal gyrus | L | 1176 | -40 | 8 | 42 | 6 |

#### 2.1.3 Data analysis

We employed the quantitative activation likelihood estimation (ALE) algorithm as implemented in the GingerALE software, v2.3.6 (Eickhoff et al., 2009, 2012; Turkeltaub et al., 2012). The ALE algorithm ultimately aims at assessing statistically whether a specific task activates each specific voxel of the brain more likely than by chance (Eickhoff et al., 2009). To account for spatial uncertainty in the location of activity, each voxel at the location of the peak-activation from each contrast and experiment is convolved with a 3-dimensional Gaussian kernel whose full-width at half maximum is weighted by the number of subjects used in that particular experiment. The sum of these Gaussians constitute the modeled activation (MA) maps. We used a within-cluster p < 0.05 FWE correction with p < 0.001 uncorrected as the cluster-forming threshold. The minimal cluster size was set to 200mm^3^. Visualization of foci and the activation clusters from the above mentioned analyses was performed using MRIcron software (http://people.cas.sc.edu/rorden/mricron/index.html) and the clusters were labeled using the Automated Anatomical Labeling (AAL) atlas (Tzourio-Mazoyer et al., 2002) and Brodmann atlas implemented within the same software.

### 2.2. Experimental and clinical studies on bodily self-consciousness (BSC)

As there are only very few published studies on global aspects of BSC including changes in first-person perspective (see Blanke et al., 2015; Grivaz et al., 2017) we included data from our previously performed fMRI study in twenty-two healthy subjects (M = 25.4, SEM = 5.7, 22 male), assessing neural mechanisms of BSC using multisensory stimulation. In detail, Ionta et al. (2011) experimentally manipulated two global aspects of BSC (i.e. self-location and first-person perspective) by using an MRI-compatible robot. The BSC regions from the fMRI study were located at the left and right temporo-parietal junction (TPJ) and included the posterior part of the superior temporal gyrus, the parietal operculum, the posterior insula and superior portion of the supramarginal gyrus (lTPJ MNI: −54, −32, 20; rTPJ MNI: 55, −28, 16). We also included results from our previously performed lesion analysis on nine neurological patients, suffering from altered states of BSC (characterized by an abnormal self-location and first-person perspective) caused by brain damage (Ionta et al., 2011). The quantitative lesion analysis of these patient data revealed a maximal lesion overlap at the rTPJ and included the posterior end of the superior and middle temporal and angular gyri (MNI: 54, −52, 26), which was located somewhat posterior to the fMRI-based BSC regions. The union of the regions from the fMRI and lesion analyses defined our BSC regions **(Table 1, Figure 1A)**.

**Figure 1.**
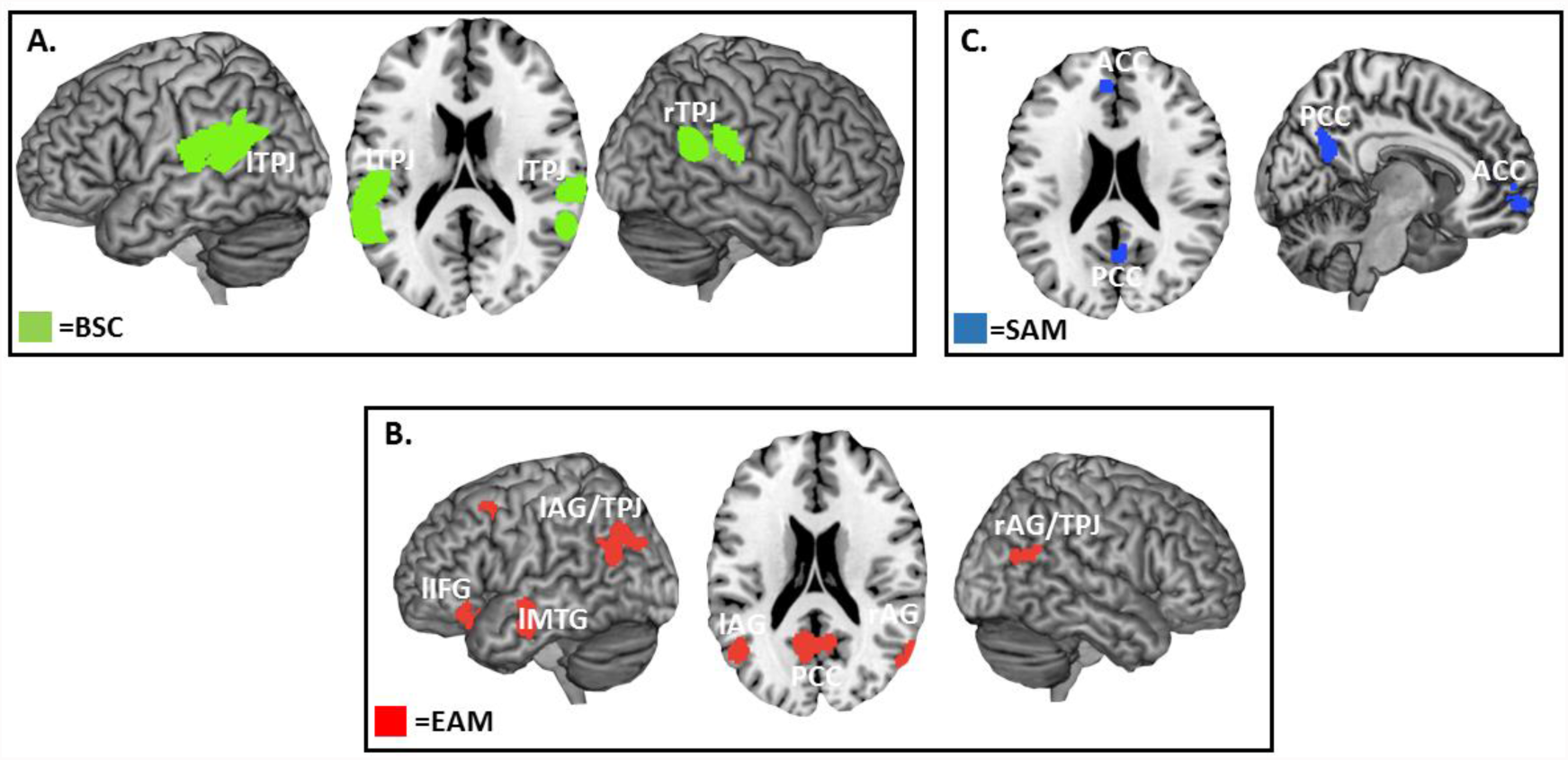
ALE-results for BSC, EAM, and SAM. **Figure** **1A**. Visualization of the BSC (in green; i.e. self-location and first-person perspective) results. **Figure** **1B**. Visualization of the parietal regions of the EAM ALE analysis results (in blue). **Figure** **1C**. Visualization of the cortical midline structures of the SAM ALE analysis (in red). Within-cluster FWE-corrected p < 0.05 with p < 0.001 (uncorrected) as the cluster forming threshold. Label provided using the MRIcron. TPJ: temporal-parietal junction; ACC: anterior cingulate cortex; PCC: posterior cingulate cortex; AG: angular gyrus; IFG: inferior frontal gyrus; MTG: middle temporal gyrus; SFG: superior frontal gyrus.

## 3. Results

### 3.1.1. EAM regions (individual ALE analysis)

The ALE meta-analysis of the forty-one EAM studies uncovered ten clusters **(Table 2, Figure 1B**). More specifically, we found activation bilaterally in the angular gyrus (AG, Brodmann area 39) and the left superior temporal gyrus (STG, Brodmann area 22). We found consistent activations in cortical midline structures, i.e. bilaterally in the ventral and dorsal parts of the posterior cingulate cortex (PCC; Brodmann areas 31 and 23) and left anterior cingulate cortex (ACC, Brodmann area 10). The analysis also revealed activity bilaterally in the parahippocampal gyri (PHG; Brodmann areas 28, 35, 36) and in the left inferior temporal gyrus (ITG, Brodmann area 21). Other clusters were found in the left inferior frontal gyrus (IFG, Brodmann area 47), bilaterally in superior frontal gyrus (SFG, Brodmann area 32) and in the left middle frontal gyrus (MFG, Brodmann area 6).

### 3.1.2. SAM regions (individual ALE analysis)

The ALE meta-analysis of the twenty-five SAM studies uncovered two clusters **(Table 3, Figure 1C)**. These were located in the cortical midline structures, i.e. bilateral ventral and dorsal parts of the ACC (Brodmann areas 32, 24) and bilaterally in the ventral and dorsal PCC (Brodmann area 31 and 23). There was no activity in the region of the angular gyri.

**Table 3.**
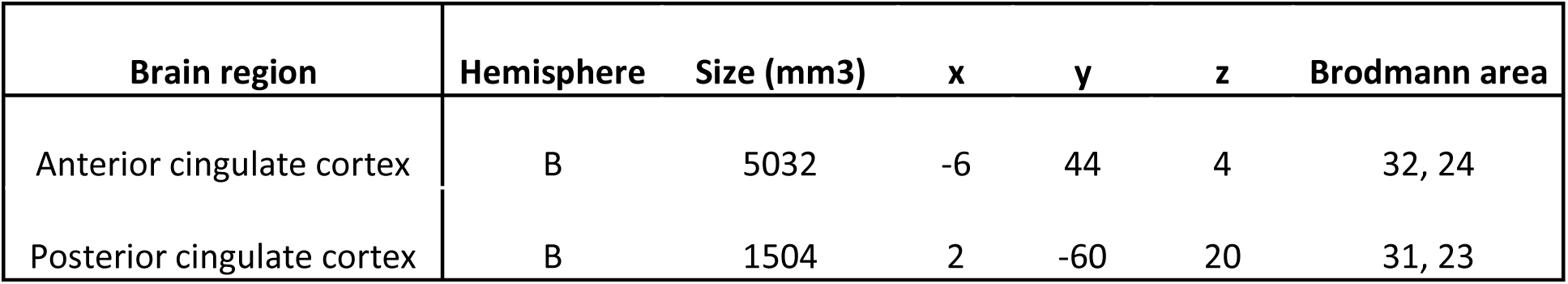
Results from SAM meta-analysis. Within-cluster FWE-corrected p < 0.05 with p < 0.001 (uncorrected) as the cluster forming threshold. R= right, L= left, B= bilateral; x, y, z are the coordinates in MNI space. Label provided using the Brodmann atlas *active cluster size in voxels

### 3.1.3. Anatomical overlap between BSC, EAM and SAM

We found an anatomical overlap between EAM regions and BSC regions bilaterally in the angular gyrus (Brodmann area 39) **(Figure 2A)**. EAM and SAM overlapped in the ventral and dorsal parts of the PCC (Brodmann areas 31 and 23) and bilaterally in the dorsal part of ACC (Brodmann area 32) **(Figure 2B)**.

**Figure 2.**
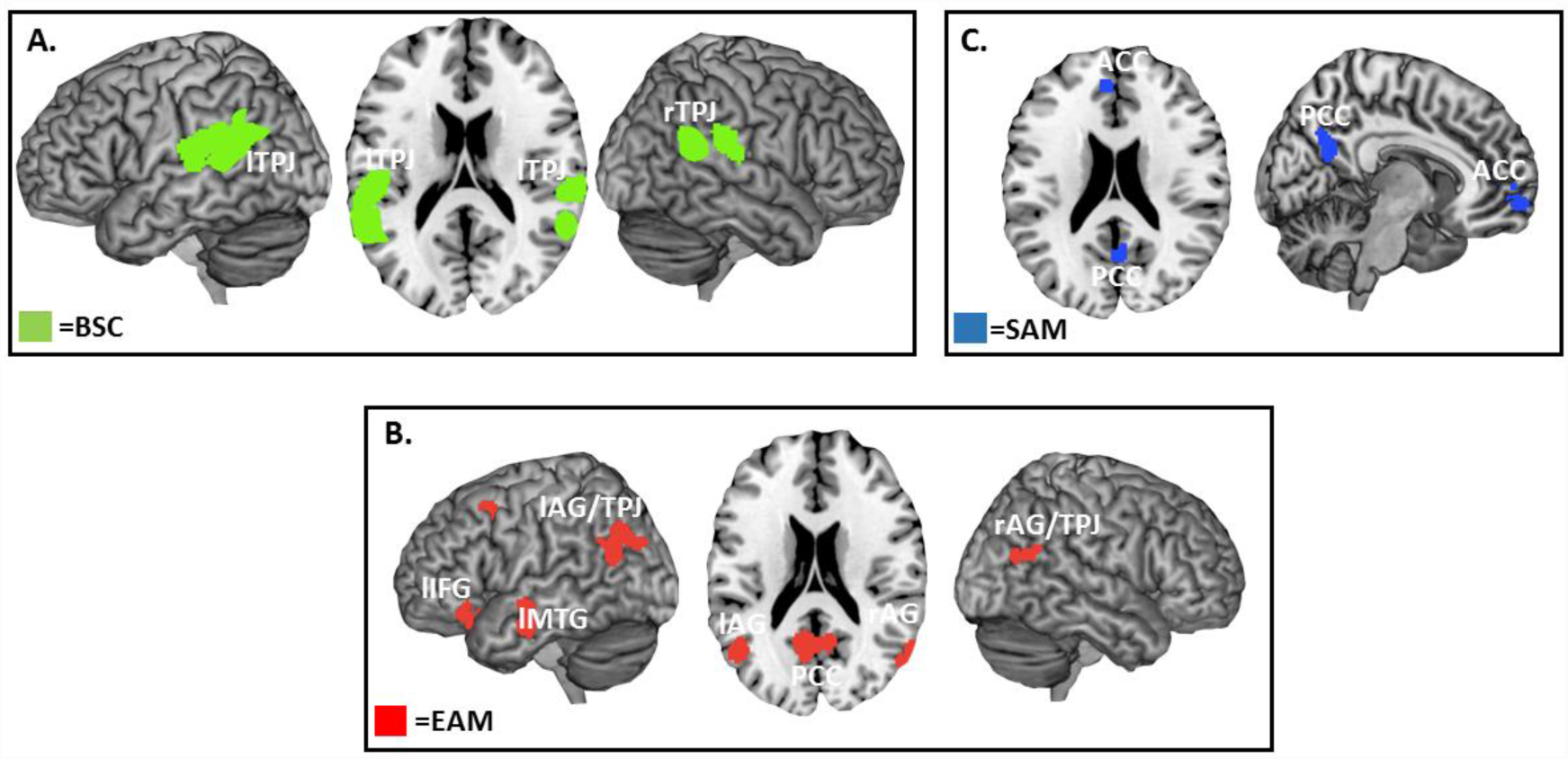
Results of overlap analysis. **Figure** **2A**. Visualization of the anatomical overlap between BSC (Ionta et al., 2011) and EAM ALE analysis in bilateral parietal cortex. **Figure** **2B**. Visualization of the anatomical overlap between SAM and EAM ALE analyses in the cortical midline structures. Within-cluster FWE-corrected p < 0.05 with p < 0.001 (uncorrected) as the cluster forming threshold. Label provided using the MRIcron.

## 4. Discussion

The current analysis revealed bilateral IPL activations in the AG (Brodmann area 39) for EAM, but not SAM. This finding is in line with previous neuropsychological and neuroimaging studies (Addis et al., 2016; Foster et al., 2015; Kim, 2016; Kim et al., 2012; Svoboda et al., 2006), consistently reporting the IPL region as the second most frequently activated region outside MTL structures during EAM retrieval. Although IPL has been often associated with visuospatial perception, attention (Corbetta and Shulman, 2002), and multisensory integration (Driver and Noesselt, 2008; Tomasino and Gremese, 2016), it is only more recently that studies associated the IPL and especially the AG with retrieval related to EAM (Bellana et al., 2017; Cabeza et al., 2007; Gilmore et al., 2015; Igelström and Graziano, 2017; Moscovitch et al., 2016; Rugg and Vilberg, 2013; Rutishauser et al., 2017; Sestieri et al., 2011, 2017; Wagner et al., 2005). This has led to several suggestions about the contributions of the parietal cortex, especially AG, to EAM retrieval. For example, the working buffer hypothesis (Vilberg and Rugg, 2008) proposes different roles for the ventral and dorsal regions in lateral parietal cortex, linking ventral parietal cortex to EAM recollection (i.e. the ability to subjectively re-experience past events enabled by autonoetic consciousness) and the dorsal parietal cortex to familiarity related processing (i.e. general recognition of events without any details associated with noetic consciousness) (Cabeza et al., 2008). Recently, Bonnici et al. (2016) investigated the role of the AG during retrieval of unimodal and multimodal episodic and semantic memories. Their findings suggest that the AG may enable the multimodal (i.e. audio-visual) integration of sensory features into rich, vivid and subjectively relevant EAM, although the precise role of AG in memory and autobiographical memory is still a matter of controversy. The present ALE data on EAM corroborate these proposals about the IPL’s involvement, including the AG and adjacent regions of the superior temporal gyrus in EAM, but not in SAM, as tested by the included studies.

Our findings further show that EAM activity anatomically overlaps with BSC, as studied by Ionta et al (2011), bilaterally in the AG. The present study extends previous EAM investigations by investigating the role of self-related processing in autobiographical memory, based on spatial aspects of BSC (self-location and first-person perspective). Previous work on BSC investigated self-related processing by using multisensory cues through a variety of experimental paradigms, highlighting contributions from own body signals (Bergouignan et al., 2014; Blanke, 2012; Blanke and Metzinger, 2009; Blanke et al., 2015; Ehrsson, 2007; Serino et al., 2013; Tsakiris et al., 2010). Given the link of BSC with subjective experience and seminal proposals by Endel Tulving that subjective re-experiencing of specific past events is a fundamental component of EAM (Tulving, 1985b, 2005), we speculated that neural processes related to multisensory bodily processing are not only relevant for BSC, but also for consciousness concerning past events and EAM (i.e. autonoetic consciousness). During the encoding of personal-life episodes, signals that are of relevance for BSC are always perceptually present, albeit in the background in most instances. For instance, Bergouignan et al. (2014) reported that recall of EAM items and hippocampal activity during the encoding of episodic events is modulated by the visual perspective from where the event was viewed during encoding. Using different methodology, St. Jacques et al. (2017) showed that first-versus third-person perspective during memory retrieval modulated recall of autobiographical events, associated this with medial and lateral parietal activations. In a recent study Bréchet et al. (2018), VR technology has allowed us to experimentally control and manipulate key elements of BSC during EAM encoding and retrieval, revealing that classical BSC factors also influence EAM performance. Our current findings are compatible with these data on shared resources for BSC and EAM and reveal an anatomical overlap bilaterally at the level of AG. As expected, we found no overlap between spatial aspects of BSC and SAM.

More work is needed given that we focused our analysis on activations reported by Ionta et al. (2011) and because distinct aspects of BSC (i.e. Grivaz et al., 2017; Guterstam et al., 2015) and other aspects of self-related processing (i.e. Qin and Northoff, 2011) have been shown to recruit cortical midline structures, close to the two overlap SAM regions revealed in the present study. A limiting factor of the present study is that there are currently only very few studies on BSC and even fewer on experimentally altered states of spatial aspects of BSC (self-location and first-person perspective) that are of particular relevance for EAM. More experimental work is needed on such spatial aspects of BSC to extend the present findings to meta-analytic work on such BSC aspects. We note that other aspects of BSC, such as full-body ownership, recruit at least partly different brain regions and may overlap in different EAM regions, including distinct parietal lobe regions as those reported here (for review see Blanke et al., 2015; Grivaz et al., 2017) and may involve MTL structures (i.e. Bergouignan et al., 2014).

Concerning the IPL, we here demonstrated that AG, but not SMG or adjacent parts of the superior temporal gyrus (revealed by the study of Ionta et al. (2011)) were jointly involved in BSC and EAM. Unlike neurological patients with damage to MTL structures (Scoville, 1957; Steinvorth et al., 2005), patients with lateral parietal lobe damage do not suffer from severe EAM deficits. For this reason, the subtler EAM impairments associated with damage to the IPL were previously largely overlooked. However, recent clinical studies investigating memory in patients with parietal lesions have provided valuable insights and especially highlighted contributions of the AG to EAM (Ben-Zvi et al., 2015; Berryhill et al., 2007; Hower et al., 2014; Levy, 2012; Simons et al., 2010). Even though patients with damage to parietal cortex are able to retrieve past personal events, a number of such lesion studies revealed that ventral IPL, especially the left AG, is associated with the impairment of subjective re-experiencing of vivid, rich and multi-sensory EAM. Moreover, patients with parietal lobe damage often report lower confidence in the retrieval of their EAM. These findings suggest a particular role of AG in the subjective experience of EAM recollection and indicate that the MTL may not be sufficient for the full-blown subjective experience of the self in the past. Based on the present meta-analytical findings we suggest that BSC-related processing in bilateral AG (with lesser or no involvement of SMG and superior temporal gyrus) may play an important role in EAM, especially with respect to subjective re-experiencing of EAM.

## Conclusion

Recent data in cognitive neuroscience suggest that the IPL contributes to EAM retrieval. The present study extends earlier neuroimaging and neuropsychological patient work and provides evidence for shared neural parietal resources between BSC and EAM, but not SAM. Our data suggest that conscious re-experiencing of past life episodes (EAM) and the conscious on-line experience of being located and experiencing the world first-person (BSC) both depend on IPL structures. We find that especially the AG - and not the SMG or adjacent parts of the superior temporal gyrus - is jointly involved bilaterally in processing related to BSC and EAM and may be an important structure for neural processing related to self-consciousness including conscious online experiencing and later re-experiencing in EAM.

---

**Supplementary Table 1.** Overview of studies investigating neural correlates of episodic autobiographical memory (EAM).

fMRI= functional magnetic resonance

PET= positron emission tomography

**Supplementary Table 2.** Overview of studies investigating neural correlates of semantic autobiographical memory (SAM).

fMRI= functional magnetic resonance

PET= positron emission tomography

